# Worldwide strains of the nematophagous fungus *Pochonia chlamydosporia* are endophytic in banana roots and promote plant growth

**DOI:** 10.1101/2020.06.10.144550

**Authors:** Cristina Mingot-Ureta, Federico Lopez-Moya, Luis Vicente Lopez-Llorca

## Abstract

The biocontrol fungus, *Pochonia chlamydosporia*, colonizes endophytically banana roots. Root hairs and root surface were found colonize by the fungus using a stable GFP transformant. Hyphal penetration of root cells was also observed. Spores of *P. chlamydosporia* 123, significantly increase root and leaf length and weight in banana plantlets *(Musa acuminata* cv. ‘Dwarf Cavendish’) in growth chamber experiments 30 days post-inoculation (dpi). In greenhouse 8L pot experiments, *P. chlamydosporia* 123 spore inoculation significantly increases leaf and root length and leaf weight in banana plants (75 dpi). Spore inoculation of *P. chlamydosporia* strains from worldwide origin (Pc21 Italy, Pc123 Spain, Pc399 China, and Pccat Cuba), significantly increases root, corm and leaf length and weight in banana plantlets. Pc21 was the best colonizer of banana roots. Consequently, this strain significantly increases most banana root and leaf length. Root colonization by *P. chlamydosporia* was also detected using cultural techniques and qPCR.

## 1. INTRODUCTION

Banana *(Musa acuminata)* is a monocotyledonous plant belonging to the Musaceae family. It is a large herbaceous plant, but does not possess a true trunk, rather foliar pods that form the pseudostem (Gold et al., 2001). Banana fruit is considered the fourth most important crop in the world (Perrier et al., 2011). Crop yield is conditioned by a set of biotic and abiotic factors, whereby many of the biotic interactions (positive or negative) that influence crops occur in the rhizosphere (Pingali, 2012). Many important pest and diseases can affect banana plants, such as black weevil (*Cosmopolites sordidus*), plant pathogenic fungi (*Fusarium oxysporum* f. sp. cubense) and plant-parasitic nematodes (Dubois et al., 2004). Nematodes are plant pathogens that inhabit agricultural crop soils reducing crop yield and causing significant economic losses in banana plantations (Davies & Elling, 2015; dos Santos et al., 2017). Biological control agents can be used for the safe management of plant-parasitic nematodes such as *Meloidogyne* spp. (Manzanilla-Lopez et al., 2013; Escudero et al., 2016). *Pochonia chlamydosporia* (Pc) (Gams & Zare, 2001) is a root-knot nematode egg-parasite (dos Santos et al., 2013; Giné et al., 2013). This fungus has been studied extensively for sustainable management of nematodes affecting crops of economic importance such as cucumber (Viggiano et al., 2014). An application of 5,000 chlamydospores per gram of soil can reduce damage caused by *M. incognita* in tomato (Atkins et al., 2003; Yang et al., 2013). This fungus is also a saprophyte in the soil, influenced by root exudates (Ward et al., 2012; Escudero et al., 2014). Pc is a true root endophyte able to prime plant root cells (Bordallo et al., 2002; Zavala et al., 2017). During plant colonization Pc obtains nutrients and protection from the host plant (Cheng et al., 2019). Pc enhances plant growth due to the production of secondary metabolites such as indole-3-acetic acid (IAA), cytokinins, gibberellic acid, ethylene and other plant growth metabolites (Bordallo et al., 2002; Tsavkelova et al., 2006; Zavala et al., 2015; Larriba et al., 2015; Zavala et al., 2017). This fungus can confer beneficial effects on mono and dicot plants (Bordallo et al., 2002; Macia-Vicent et al., 2009a). All these properties make this fungus a good candidate for banana plant biomanagement.

In this work, we devise a new method for inoculating Pc123 in banana plants. We also test worldwide Pc strains to determine the effect of fungus genetic variation in plant growth promotion and root colonization.

## 2. MATERIALS AND METHODS

### 2.1 Plants and fungi

*In vitro* 30-day-old banana plantlets (*Musa acuminata* cv. ‘Dwarf Cavendish’ AAA) were obtained from Cultesa (Santa Cruz de Tenerife, Canary Islands, Spain). Before experiments, plantlets were acclimatized in a growth chamber (SANYO MLR-351H) for 5-10 days, at 24 °C, 60% relative humidity and 16h/8h light/darkness photoperiod.

Pc21, Pc123, Pc399 and Pccat strains from the nematophagous fungus *Pochonia chlamydosporia* (Pc) were obtained from the Collection of the Phytopathology Laboratory (University of Alicante, Spain). Pc21 was isolated in Metaponto (Italy) from *Meloidogyne* sp. in kiwi trees (Macia-Vicent et al., 2009a) and was a kind gift from Dr. A. Ciancio (CNR, Bari, Italy). Pc123 was isolated from *Heterodera avenae* eggs in southwest Spain (Olivares-Bernabeu & Lopez-Llorca, 2002). Pc399 was obtained in China by late Professor Brian Kerry (Rothamsted Research, UK), who kindly sent us the strain for research purposes. Pccat was isolated from *Meloidogyne spp*. infected eggs in Cuba (Zavala et al., 2015) and was a kind gift from Dr. L. Hidalgo (CENSA, Cuba).

### 2.2 Growth chamber experiments

Plantlets were grown individually in 200 ml polystyrene cups with sterilized Terraplant 2 peat (Compo Expert, Castellón, Spain). The experiment included three treatments with Pc123 and control. Treatment 1 consisted of plantlets inoculated with eight 5mm diameter cores from the edge of a 21-day-old Pc123 colony on CMA. In Treatment 2, plantlets had roots placed on Pc123 colonies for 5 days in Magenta boxes (GA-7, Sigma) with CMA. These plantlets were then transferred individually to polystyrene cups with peat. In Treatment 3, plantlets were inoculated with 10,000 Pc123 conidia and chlamydospores (c/c) per gram of substrate. Controls were polystyrene cups with sterilized peat and a banana plantlet only. Plantlets were irrigated every 48h with sterilized 1/10 Gamborg’s salt solution (Basal Medium Minimal Organic, Sigma). Control and treated plantlets were grown in a growth chamber as described (see 2.1). Thirty days after planting, root, corm and leaf weight and length per plant were assessed. The number of leaves per plant was also recorded. Root colonization by Pc was estimated using culturing techniques (see 2.5). This experiment was repeated three times with 10 plantlets per treatment in each replicate.

To visualized Pc development in roots, banana plantlets were inoculated in Magenta boxes with a Pc 123-GFP transformant strain (Macia-Vicente et al., 2012). Seven days after inoculation, roots were sampled. Root fragments were then excited with a 488 nm laser, and GFP fluorescence monitored at 505-530 nm with a Laser Confocal Microscope (Leica TCS-SP2).

### 2.3 Greenhouse experiments

Thirty plantlets were either inoculated with 50,000 Pc123 c/c per gram of substrate, or not (controls). Plantlets were grown in polystyrene cups, for 30 days as above (see 2.2). Five plantlets per treatment were then destructively sampled to assess root colonization by Pc using conventional PCR and qPCR techniques (see 2.5). Twenty plants per treatment were then transplanted to 8L pots and grown in the greenhouse, at 24°C and 60% relative humidity. Plants either received a single (initial) inoculation with 50,000 Pc123 c/c (Treatment 1), or they were inoculated monthly (three times) with this same inoculum (Treatment 2). Control plants were left uninoculated. Plants were irrigated daily with 100 ml tap water and were grown for 75 days. Root, corm, pseudostem and leaf weight and length per plant were then scored. Number of leaves per plant was also determined. Root colonization by Pc was estimated using culturing and PCR techniques (see 2.5). This experiment was repeated twice with 20 plants per treatment in each replicate.

### 2.4 Pc strains variability and banana growth promotion experiments

We used Pc21, Pc123, Pc399, and Pccat in banana plantlets as described above (see 2.2). The experiment design and conditions were the same as above (see 2.2). Treatments consisted of a single initial inoculation of 50,000 c/c from a given Pc strain per treatment. Thirty-day-old plantlets were scored as above (see 2.2). Pc root colonization was also determined using culturing and PCR techniques (see 2.5).

### 2.5 Root colonization

Ten root systems per treatment from plants of either growth chamber or greenhouse experiments were sampled to evaluate root colonization by Pc. Five root systems were used for total root colonization and the rest for endophytic colonization estimations. For total root colonization, roots were rinsed in sterile distilled water three times for one minute each. For endophytic colonization, a preliminary 1min wash in 1% sodium hypochlorite was performed. From each root system, twelve 1 cm pieces were randomly sampled axenically. Root samples were then plated on a growth restricting medium (Lopez-Llorca & Duncan, 1986). Petri plates with root fragments were incubated for 20 days at 24°C in darkness.

Alternatively, root fragments were also employed to measure root colonization by Pc with PCR using specific primers (Table 1) (Escudero & Lopez-Llorca, 2012). An improved cetyl-trimethyl-ammonium bromide (CTAB) extraction method was used for isolating genomic DNA from lyophilized samples (O’Donnell et al., 1998; Lopez-Llorca et al., 2010). PCR was performed using 100ng DNA mixed with Routine Polymerase 2X (Thermo Fisher) and with each primer at 0.5 μM. Negative controls contained sterile water only, and positive ones contained 100ng/μl of Pc123 DNA extracted from axenic cultures. DNA amplification was carried out in PTC-100 Peltier Thermal Cycler (Escudero & Lopez-Llorca, 2012). PCR products were run in 1.5% agarose gel electrophoresis stained with GelRed (Biotium, Hayward, USA).

**Table 1.**
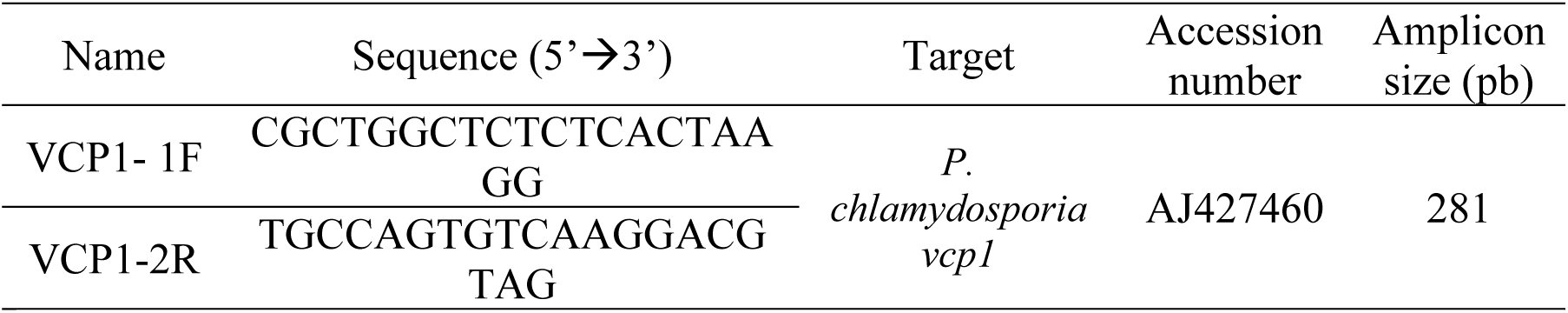
Primers used to detect/ quantify the fungus *Pochonia chlamydosporia* in banana plant roots.

Quantitative PCR (qPCR) was performed in triplicate using a 96-well plate-based (Roche Diagnostics, Penzberg, Germany) in a Thermal Cycling StepOnePlus (Applied Biosystems, Foster City, CA, USA), as described in Escudero & Lopez-Llorca, (2012)). Root DNA extracts (10ng) from each growth chamber treatment were mixed with FastStart Universal SYBR Green Master (Roche, Barcelona, Spain) and with each primer at 0.5 μM. Negative controls contained sterile water. Standard curves were produced by plotting the average cycle threshold (Ct) values against the log DNA concentration, amplification efficiencies were calculated from the slope of Ct plotted against log DNA concentration (Macia-Vicente et al., 2009b; Escudero & Lopez-Llorca, 2012).

### 2.7 Statistical analysis

The normality of the data was verified by the Shapiro-Wilk test, and the Levene test was used to study the homogeneity of the variance between the groups. Data that followed a normal distribution were compared using a two-way ANOVA for differences between treatments and/or sampling times. Non-normal data were compared using the Kruskal-Wallis test (K-W). Wilcoxon test was used for multiple tests for pairwise comparisons. In all cases, the level of significance considered was 95%. All analyses were performed using the R software (V.3.4.3) (R Core Team 2017) and GraphPad Prism version 8 for Windows (GraphPad Software, Inc., San Diego, CA, EE. UU.).

## 3. RESULTS

### 3.1 Pc spore inoculation is the most effective inoculant for promoting plant growth

*P. chlamydosporia* (Pc123) conidia and chlamydospores significantly promoted root and leaf growth of banana plantlets (Fig. 1a,b). Pc123 can colonize endophytically banana roots using both, mycelium and spores as inoculum in plantlets at 10-20-30 dpi (Fig 1c). Root colonization by Pc123 can be detected from 10 dpi onwards using all treatments. We also detected Pc123 GFP transformant in the rhizoplane and root cells of *M. acuminata* 7dpi (Fig 1d,e). Roots inoculated with conidia and chlamydospores show at 30 dpi higher colonization by Pc123 than other treatments evaluated. Therefore, conidia and chlamydospore were used as the inoculum for further experiments.

**Figure 1.**
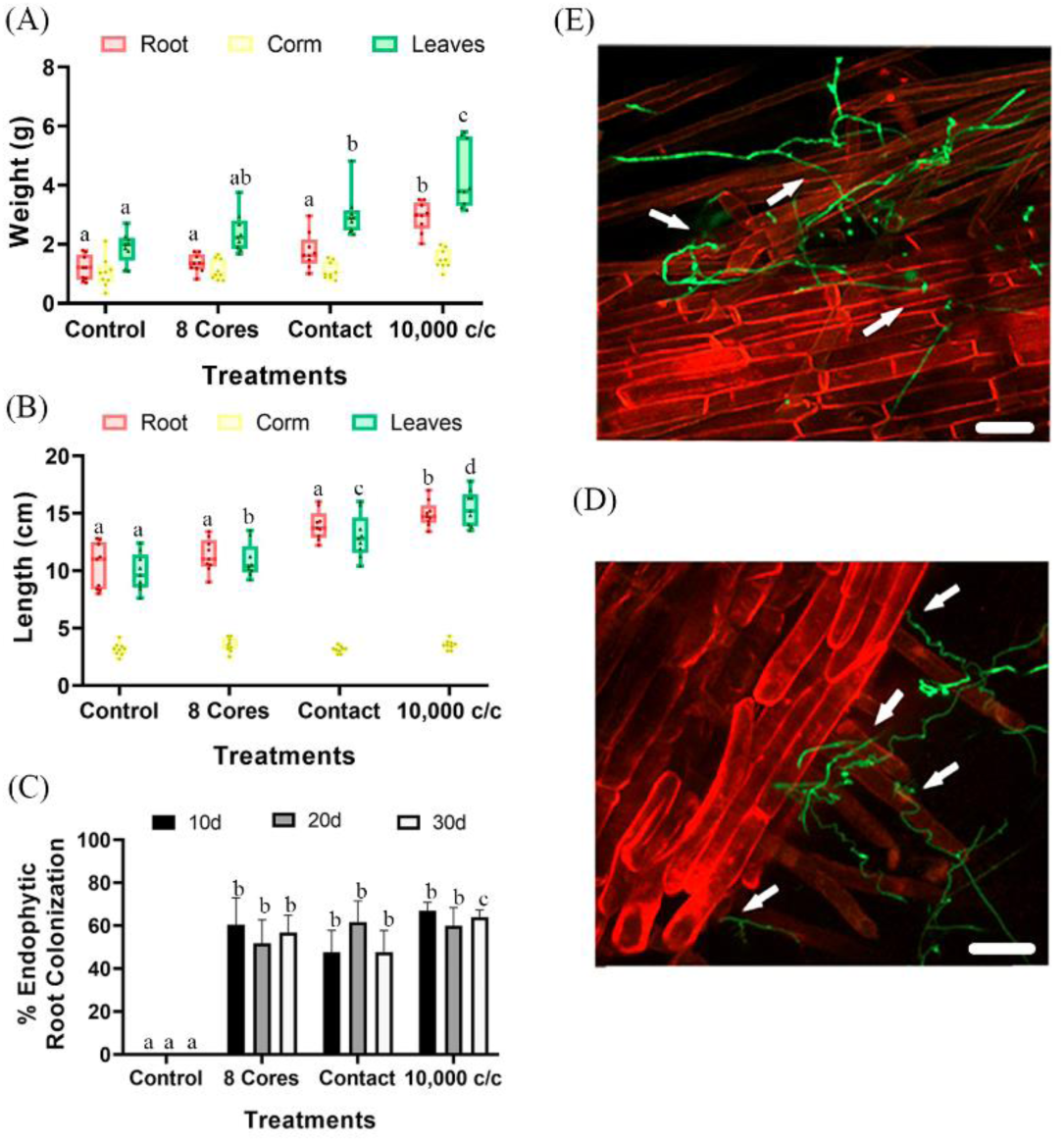
Effect of *P. chlamydosporia* on banana plantlets. Treatments: (8 cores) from the edge of 21-day-old Pc colony; (Contact) of banana plantlet in Magenta Box with Pc for 5 days; (10,000c/c) suspension of Pc; (Control) plantlets without fungal inoculation. (a) Maximum Root, Leaf and Corm Length. (b) Fresh Root, Leaf and Corm Weight. (c) Total and Endophytic root colonization by *P. chlamydosporia*. In the parameters analyzed individually, treatments with different letters indicate significant differences (p-value <α = 0.05). (d) Root hairs colonization; (e) Rhizoplane colonization of *M. acuminata* by Pc123 GFP modified strain. Arrows: indicate possible hyphal penetration points in the root. White bar=75 μm.

### 3.2 Pc promotes the growth of banana plants in the greenhouse

*P. chlamydosporia* (Pc123) significantly increases root and leaf length of banana plants (Fig. 2a) under greenhouse conditions. Pc123 also increases leaf weight of banana plants in all treatments (Fig. 2b). After 30d we checked that Pc was well established in banana plant roots. Root colonization was verified by PCR (Fig. 2c) and qPCR (Fig. 2d). Root colonization by Pc in older banana plants (75dpi) was estimated by culturing and PCR (Fig. 2e,f). qPCR standard curve shows a R^2^ higher than 0.89 (Fig. S3). Molecular quantification of Pc123 in banana root, revealed more Pc123 (approx. 3-fold) in total *vs*. endophytic colonization (Fig 2c). Pc123 can persist endophytically in banana roots up to 75 dpi (Fig. 2e). Root colonization by Pc123 increases with repeated applications of the fungus.

**Figure 2.**
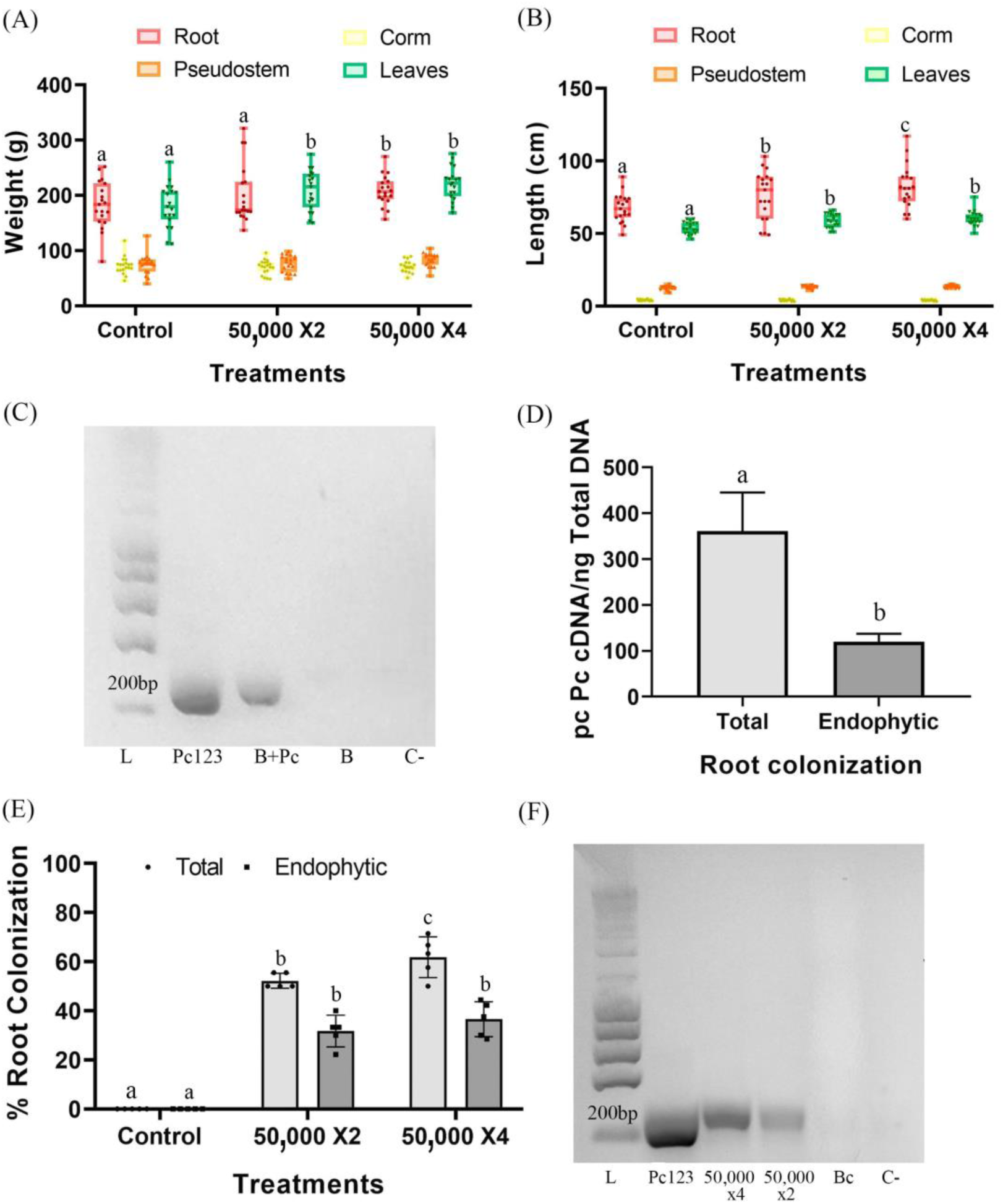
Effect of *P. chlamydosporia* 123 on the growth of banana plants. Treatments: (50,000 x2) two spore inoculation plants; (50,000 x4) four spore inoculation plants; (Control) plants without fungal inoculation. (a) Maximum Root, Leaf, Pseudostem and Corm Length. (b) Fresh Root, Leaf, Pseudostem and Corm Weight. (c) Total and Endophytic Root Colonization by Pc. In the parameters analyzed individually, treatments with different letters indicate significant differences (p-value <α = 0.05). (d) Quantification by qPCR of *P. chlamydosporia* colonization of banana roots in 30days old plants. (e) Molecular detection of *M. acuminata* root colonization by *P. chlamydosporia*., 30 dpi plants. (f) Molecular detection of *M. acuminata* root colonization by *P. chlamydosporia*. In 105-days-old plants. PCR of the *P. chlamydosporia vpc1* gene. Abbreviations: (M) Ladder; (Pc) DNA extracted from the mycelium of *P. chlamydosporia*; (B + Pc) DNA extracted from 30 dpi roots of the plant (B) inoculated by *P. chlamydosporia* (Pc); (B) DNA extracted from the root of a 30-days banana plant without inoculation; (50,000 x4) DNA extracted from four spore inoculation 105 days old plants; (50,000 x2) DNA extracted from two spore inoculation 105 days old plants; (Bc) DNA extracted from the root of a 105-day-old banana plant without inoculation; (C-) negative control without DNA.

### 3.3 Pc strains vary in their plant growth promotion ability

All Pc strains tested caused a significant increase in weight and length in banana plantlets (Fig. 3a,b). Pc21 was the most banana growth promoting strain. Banana plantlets inoculated with Pc123, Pc399 and Pccat also show growth promotion. Pc21 was the largest colonizer of banana roots (total colonization) (Fig. 3c). Primers designed from the Pc123 *vcp1* gene sequence could amplify the gene from other apart from that of Pc123 (Fig. S5.) These strains were detected in banana roots 30 dpi. As seen in Figure 3d, there was root colonization by all Pc strains.

**Figure 3.**
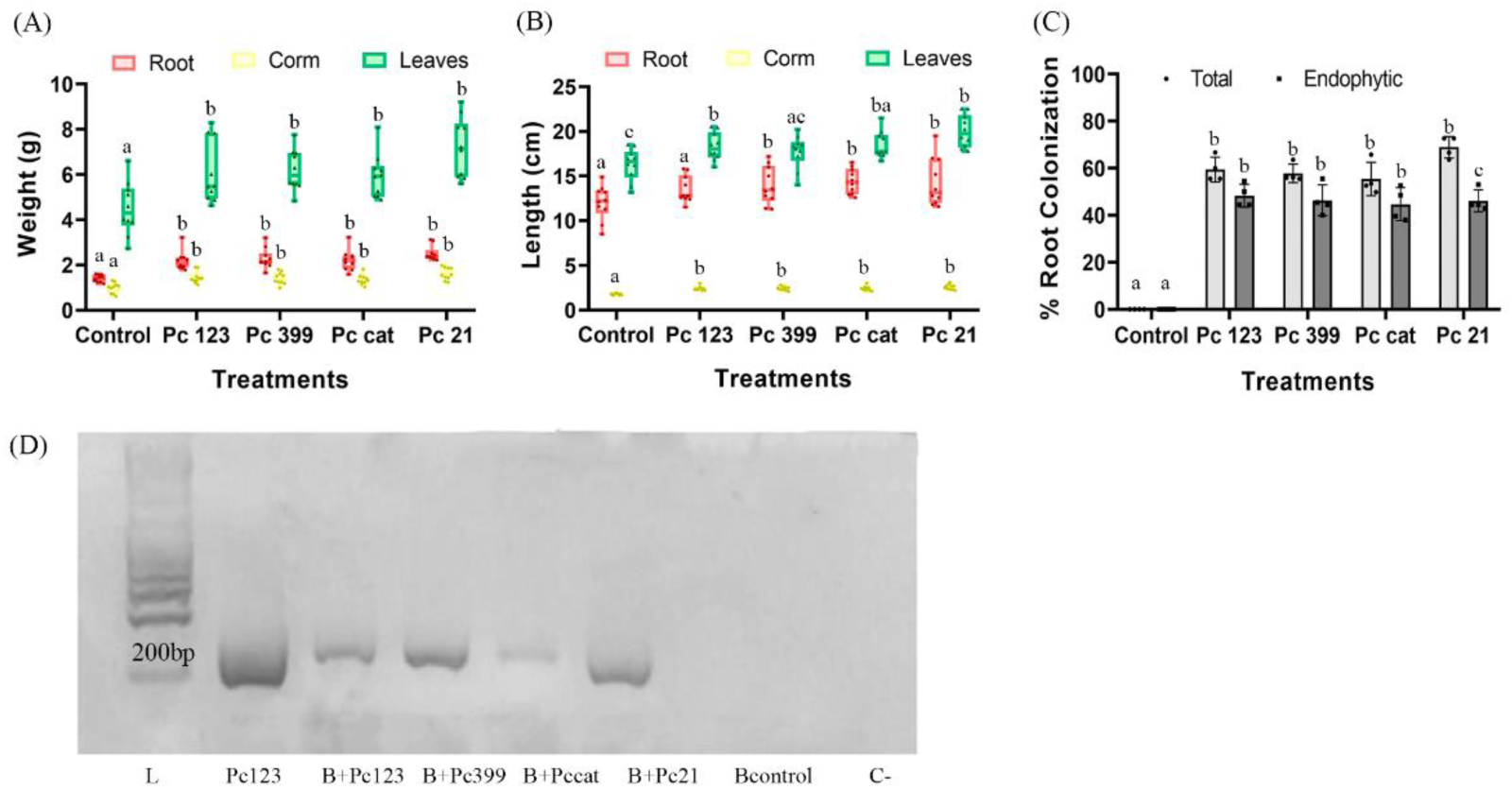
Effect of *P. chlamydosporia* diversity on the growth of 30-day-old plantlets. Treatments: inoculation of 50,000 conidia and chlamydospores suspension of each Pc strains (Pc123, Pc399, Pccat and Pc21). (a) Root, Leaf and Corm Length. (b) Fresh Root, Leaf and Corm Weight. (c) Total and Endophytic Root Colonization. Growth indicators analyzed individually, the treatments with different letters show significant differences (p-value <α = 0.05). (d) Molecular detection of *P. chlamydosporia* strains colonizing M. *acuminata* roots, using the *vpc1* gene. Abbreviations: (M) Ladder ; (B+Pc123, B+Pc399, B+Pc cat, B+Pc21) DNA extracted from 30 dpi roots of banana plant with different strains of *P. chlamydosporia*; (Bcontrol) DNA extracted from 30-day plant *(M. acuminata)* roots without *P. chlamydosporia*; (C-) negative control without DNA.

## 4. DISCUSSION

The banana crop is paramount for food security and therefore it is the most consumed and cultivated fruit worldwide (Ferri et al., 2012). In view of the burst of human population in the last 50 years, the increase in food production requires improvement of crop management (Pingali, 2012). Traditionally chemical control has been used to enhance banana yield worldwide. Phytochemicals can infiltrate in water and this has been associated with human chronic neurobehavioral dysfunctions (Wesseling et al., 2002; Diepens et al., 2014). To reduce these impacts, sustainable practices for banana crop protection are investigated (PROMUSA, 2018). Genetic modification for banana improvement has been used for biotic and abiotic stress tolerance (Ghag & Ganapathi, 2017). These authors discussed that defence peptides expressed in banana imparted resistance to pathogens but may also target beneficial fungi associated with banana roots (Backman & Sikora, 2008). To this respect, a wide diversity of root endophytic fungi colonizes wild banana plants (Zakaria et al., 2016). This microbiota is in turn modified by banana root pathogens such as *Fusarium oxysporum* (Henao et al., 2019). Ciancio et al., (2019) have recently found that the nematophagous fungus Pc is a natural component of the banana rhizosphere. This fungus is an endophyte in several crops and improves crop development and defences. In this paper, we propose the use of Pc as a banana endophyte to improve the banana crop.

We have found that Pc conidia and chlamydospores are the best inoculants for banana plant treatment, both in growth chambers and in greenhouses. To this respect, Klamic ® a Pc chlamydospore based product, was used by Hernandez-Socorro et. al. (2016) to inoculate banana *in vitro* plant AAB and AAAB cultivars. These authors found, as in our study with AAA banana, growth promotion by *P. chlamydosporia* var. catenulata (Pccat) strain. In our study, we demonstrate that Pc significantly increases root and leaf length and weight in 30-day-old banana plantlets. We have also found in greenhouses, that monthly applications of Pc spores promote root and leaf growth in three-month-old banana plants. This fungus can promote plant growth even in RKN infected banana roots (Barbosa et al., 2019). We observed that banana plant growth promotion by Pc is strain dependent. Similar results were found in tomato plants by Zavala et al. (2015), who also found strain-dependent tomato growth and yield promotion by Pc. In our study, we found Pc21 from Italy the best growth promoter in banana. The other Pc strains (Pc123, Pc399, and Pccat) increase root, leaf, and corm weight. Growth promotion by Pc has also proven in other crops such as tomato, lettuce or wheat (Monfort et al., 2005; Dallemole-Giaretta et al., 2015; Monteiro et al., 2018). This indicates that Pc is able to promote growth of a great variety of crops.

Our study establishes the basis to inoculate banana plants with Pc. So, we found spore suspensions better inoculants than mycelia treatments. Pc can be applied in combination with other biocontrol agents. Monteiro et al., (2018) found in tomato synergistic growth promotion by Pc in combination with the trapping fungus *Duddingtonia flagrans*. In our previous study (Bordallo et al. 2002) the similar trapping fungus *Arthrobotrys oligospora* decorticated mature barley roots but not tomato. In another important crop such as pea, Pc inoculation during the seed stage, also provides good results in root growth promotion (Arevalo-Ortega et al., 2019). Therefore, Pc can be inoculated at different phenological stages of the plant, enhancing plant growth and yield.

Previous studies described that Pc substantially increases growth and nutrient uptake (mainly phosphorus) in tomato plants, acting as a biofertilizer (Behie et al., 2012; Zavala et al., 2015; Monteiro et al., 2018). Phytases and phosphatases would consequently be related to growth promotion by Pc in plants (Richardson et al., 2001; Macia-Vicente et al., 2009a; Rosso et al., 2014). The fungus also synthesizes phytohormones such as (IAA), siderophores, or deaminases and other enzymes involved in plant growth (Waqas et al., 2014; Zavala et al 2015; Kumar et al., 2018; Nandhjni et al., 2018). Our results in banana growth promotion when Pc is inoculated in the rhizosphere could be related to its ability to induce IAA biosynthesis or enhancing nutrient uptake (Monteiro et al., 2018). Other studies with growth-promoting rhizobacteria found that *Bacillus* and *Pseudomonas* can also induce IAA. This phytohormone is related to root development and cellular division, promoting plant growth (Dadrasnia et al., 2020).

Using a root culturing technique (Lopez-Llorca & Duncan, 1986) we found endophytic colonization of banana roots by Pc. This fungus has been detected colonizing endophytically other plant roots, such as barley, corn, and pepper (Macia-Vicente et al., 2008; Macia-Vicente et al., 2009a; Moonjely and Bidochka, 2019). We have observed that Pc21 is the best growth-promoting Pc strain and the best root colonizer. Zavala et al., (2015) found different rhizosphere competence of Pc strains in tomato which changed with plant age. Differences in root colonization between Pc strains may be due to their abilities to modulate jasmonic acid (JA) plant response (Zavala et al., 2017). Moonjely and Bidochka, (2019) found differences in the rhizospheric competence of strains of the entomopathogenic fungus *Metarrizhium* spp. They also found Pc which was the best root colonizer with *ca*. four-fold the rhizosphere competence of their best *Metarhizium* strain. This is despite the evolutionary relatedness of both fungi (Larriba et al. 2014; Lin et al., 2018). The good rhizosphere competence of Pc in banana can be explained by the fact that Pc seems to prefer monocot to dicot plants (Moonjely and Bidochka, 2019). In our study we have also detected Pc colonization in banana roots using *vcp1*, encoding a Pc serine protease. This gene has been used to quantify Pc rhizosphere competence in previous studies (Escudero and Lopez Llorca, 2012; Zavala et al., 2015; Ghahremani et al., 2019). We have also monitored Pc colonization in banana roots by confocal microscopy using a stable GFP transformant of the fungus (Macia-Vicente et al., 2009a). The presence of Pc in roots can exclude other fungal root colonizers by competence or biosynthesis of antimicrobial compounds (Maciá-Vicente et al., 2009a; Nandhjni et al., 2018). Xue et al., (2015) found that manipulating the banana rhizosphere microbiome can control the devastating fungus *Fusarium oxysporum* f.sp. cubense tropical race 4, so Pc as endophyte could be an interesting BCA. Our study demonstrates that Pc colonizes the root system and promotes growth in *M. acuminata* plants. These results will establish the basis of Pc as a biofertilizer and bioprotectant of banana to biotic and abiotic stresses.

## ACKNOWLEDGMENTS

This research was funded by the EU H2020, Musa Project (727624). We thank members of the Laboratory of Plant Pathology (University of Alicante) for help in greenhouse experiments.

