## Supplementary material for "Worldwide strains of the nematophagous fungus *Pochonia chlamydosporia* are endophytic in banana roots and promote plant growth"

### SUPPORTING INFORMATION. FIGURE CAPTIONS

**Figure S1.** *P. chlamydosporia* effect on the number of leaves per 30-day-old banana plant. Treatments: (8 cores) from the edge of 21 days-old Pc colony; (Contact) of banana plantlet in Magenta Box with Pc for 5 days; (10,000c/c) suspension of Pc; (Control) plantlets without fungal inoculation. Treatments with different letters indicate significant differences (p-value $< \alpha = 0.05$).

**Figure S2**. Effect of *P. chlamydosporia* 123 on the number of leaves per 105-day-old banana plant. Treatments: (50,000 x2) two spore inoculation plants; (50,000 x4) four spore inoculation plants; (Control) plants without fungal inoculation. The treatments with different letters indicate significant differences (p-value $< \alpha = 0.05$).

**Figure S3.** Standard curve for real-time PCR of 4-fold serial dilutions of DNA from *P. chlamydosporia*. Cycle thresholds (Ct) were plotted against the log of known concentrations of genomic DNA standards and linear regression equations were calculated for the quantification of the unknown samples by interpolation.

**Figure S4.** Effect of *P. chlamydosporia* diversity on the number of leaves per plant. Treatments: inoculation of 50,000 conidia and chlamydospores suspension of each Pc strains (Pc21, Pc123, Pc399, and Pccat). Treatments with different letters show significant differences (p-value $< \alpha = 0.05$).

**Figure S5.** Molecular detection of *P. chlamydosporia* strains. PCR of mycelia of different strains of *P. chlamydosporia* using primers from *vpc1* gene. Abbreviations: M (ladder); Pc strains (Pc123, Pc399, Pccat, Pc21); C- (negative control, without DNA).

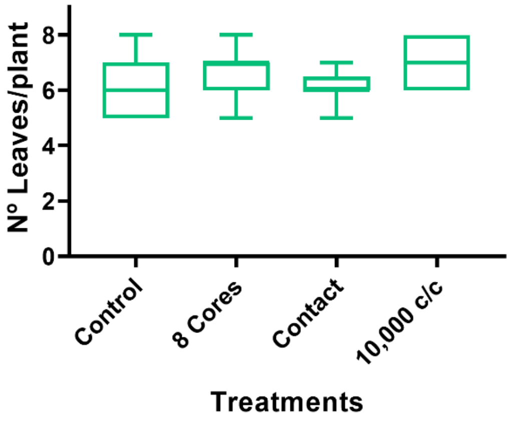

N° Leaves/plant
8
6
4
2
0
Control
8 Cores
Contact
10,000 c/c
Treatments


**Figure S1.**

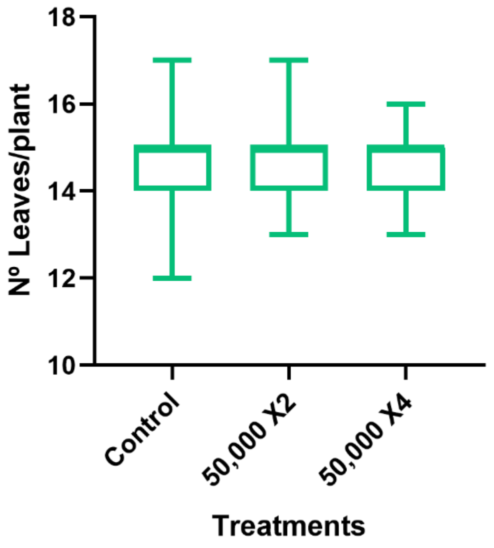
18
16
14
12
10
Nº Leaves/plant
Control
50,000 X2
50,000 X4
Treatments

**Figure S2**.

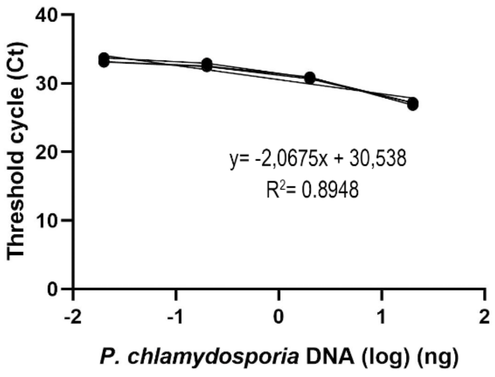
40
30
20
10
0
Threshold cycle (Ct)
y= -2,0675x + 30,538
R2= 0.8948
-2
-1
0
1
2
P. chlamydosporia DNA (log) (ng)

**Figure S3.**

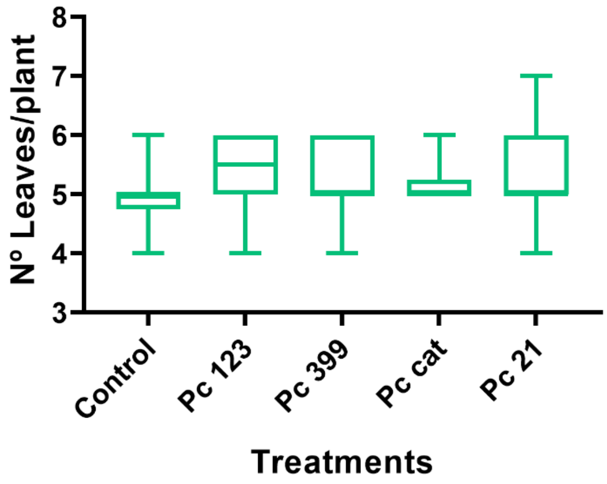
N° Leaves/plant
8
7
6
5
4
3
Control
Pc 123
Pc 399
Pc cat
Pc 21
Treatments

**Figure S4.**

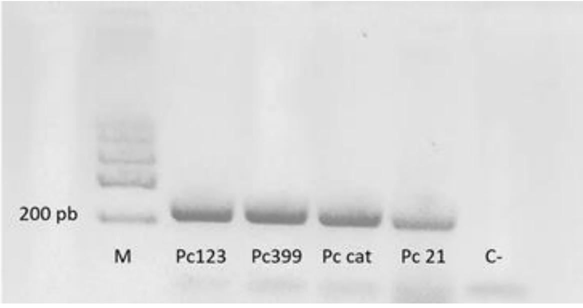


**Figure S5.**
